# M^3^CV:A Multi-subject, Multi-session, and Multi-task database for EEG-based Biometrics Challenge

**DOI:** 10.1101/2022.06.28.497624

**Authors:** Gan Huang, Zhenxing Hu, Weize Chen, Zhen Liang, Linling Li, Li Zhang, Zhiguo Zhang

## Abstract

EEG signals exhibit commonality and variability across subjects, sessions, and tasks. But most existing EEG studies focus on mean group effects (commonality) by averaging signals over trials and subjects. The substantial intra- and inter-subject variability of EEG has often been overlooked. The recently significant technological advances in machine learning, especially deep learning, have brought technological innovations to EEG signal application in many aspects, but there are still great challenges in cross-session, cross-task, and cross-subject EEG decoding. In this work, an EEG-based biometric competition based on a large-scale M^3^CV (A Multi-subject, Multi-session, and Multi-task Database for investigation of EEG Commonality and Variability) database was launched to better characterize and harness the intra- and inter-subject variability and promote the development of machine learning algorithm in this field. In the M^3^CV database, EEG signals were recorded from 106 subjects, of which 95 subjects repeated two sessions of the experiments on different days. The whole experiment consisted of 6 paradigms, including resting-state, transient-state sensory, steady-state sensory, cognitive oddball, motor execution, and steady-state sensory with selective attention with 14 types of EEG signals, 120,000 epochs. With the learning tasks of the identification and verification, the performance metrics and baseline methods were introduced in the competition. In general, the proposed M^3^CV dataset and the EEG-based biometric competition aim to provide the opportunity to develop advanced machine learning algorithms for achieving an in-depth understanding of the commonality and variability of EEG signals across subjects, sessions, and tasks.

## 1. Introduction

Since the discovery of electroencephalography (EEG) by Hans Berger in 1924 (Berger, 1929), EEG has evolved for use in a wide range of applications (Alotaiby et al., 2014; Cahn and Polich, 2006; Dietrich and Kanso, 2010; Wolpaw et al., 1991) for almost one hundred years. Typically, event-related potentials (ERPs) study focuses on the significant common or mean effects of a cohort, in which the intra- and inter-subject variability are treated as noise and are filtered out by averaging over trials and subjects (Seghier and Price, 2018). Hence ERPs have been widely used for investigating the neurological functions of sensory, motor, and cognitive processes (Kappenman et al., 2021). However, the conventional practice of ERP analysis takes the mean value of the EEG signals across trials and/or subjects to achieve a higher signal-to-noise ratio, but the group-level commonality may not have a reliable effect on the individual level (Boshra et al., 2019; Fröhner et al., 2019; Hu et al., 2021; Infantolino et al., 2018).

The intra- and inter-subject variability pose a great challenge to the individual level explanation and decoding of the EEG signal. Multiple factors may contribute to intra- and inter-subject variability, including different brain structures among subjects, non-stationarity of brain activity, neural processing for different tasks, and some unknown factors (Wei et al., 2021). With the availability and affordability of computational power, the recently significant technological advances in machine learning, especially deep learning, have brought technological innovations to EEG signals application in many aspects (Craik et al., 2019). In clinical and psychiatric studies, machine learning technique has pushed EEG-based diagnosis, prognosis, risk stratification, or treatment monitoring toward a more individualized approach (Olbrich & Conradi, 2016). In the application of brain-computer interface (BCI), intention-encoded EEG can be decoded for the direct interaction between the brain and a computer by single-trial decoding (Pfurtscheller and Neuper, 2001; Wolpaw et al., 2000). For system security, EEG-based biometrics become attractive to the research community for personal identification and verification.

However, the lack of generalizability to different subjects and different sessions limited the use of EEG-based machine learning methods in clinical and psychiatric applications (Sui et al., 2020; Fisher et al., 2018). For example, a BCI decoder usually works well in a single personal session, but performs poorly over time or even fails to be applied to another subject (Krusienski et al., 2011; Satti et al., 2010). Although there are meaningful studies concerning cross-session and cross-subject transfer learning for BCI, the theory of common feature space construction in BCI applications is still being studied (Autthasan et al., 2021; Lotte and Guan, 2011; Rodrigues et al., 2019). Similarly, severe overfitting was observed in the application of EEG-based biometrics in the within-session recognition.

In summary, the traditional ERP analysis and statistical test methods help us identify the group-level commonality of EEG, while the recently developed machine learning techniques provided us with a way to further explore the individual-level variability of EEG. However, the lack of generalizability across subjects, sessions, and tasks is still a major challenge for machine learning in neuroimaging. A large-scale multi-subject, multi-session, multi-task EEG database is highly deserved to support this branch of research. Hence, in this study, we established an M^3^CV (Multi-subject Multi-session Multi-task Commonality and Variability) EEG database to support cross-subject, cross-session, and cross-task EEG studies.

The M^3^CV database contains 14 types of EEG tasks in 6 experiment paradigms from 106 healthy young adults, in which 95 subjects completed two experimental sessions repeated on different days. To record the EEG data for as many tasks as possible within the limited recording time and to ensure the data quality of each task, we designed the experiments based on suggestions from five experts in neuroscience and psychology (see Acknowledgments).

The experimental paradigms in each session included the following six paradigms with 14 tasks of EEG signals (bolded text).

- Paradigm 1: Resting-state with eye closed and eye open (**EC** and **EO**, each lasting for 2 mins).
- Paradigm 2: Transient-state sensory with visual, auditory, and somatosensory stimulation (**VEP, AEP**, and **SEP**, each having 60 trials)
- Paradigm 3: Steady-state sensory with steady-state visual, auditory, and somatosensory stimulation (**SSVEP** 1 min, **SSAEP** 2 mins, and **SSSEP** 2 mins)
- Paradigm 4: P300 with oddball experiment (**target P300** with 30 trials, and **nontarget P300** with 570 trials)
- Paradigm 5: Motor execution with the movement of right foot, right hand, and left hand (**FT, RH**, and **LH**, each having 80 trials)
- Paradigm 6: SSVEP with selective attention (**SSVEP-SA**, six classes, each class having 12 trials, each trial lasting for 10 s)

When retrieving existing open-access EEG databases (Table I), we found few databases with multi-task and multi-session EEG signals. The number of subjects in most databases was less than 50. With some specific research objective, the number of tasks was typically only one or a small number. Multi-session EEG data are even harder to collect because it is more difficult to require all subjects to repeat the experiment after a certain time period. We also found that existing databases with some attributes of multi-task or multi-session were mainly used for reliability analysis, BCI, and EEG-based personal identification. Reliability analysis (Gaspar et al., 2011) focused on the cross-session reproducibility of EEG signals, which plays a fundamental role in EEG research. BCI studies (Brunner et al., 2008; Goldberger et al., 2000; Jeong et al., 2020; Korczowski et al., 2019; Kumar et al., 2021; Lee et al., 2019) often require the development of cross-session and cross-subject transfer learning algorithms. EEG-based personal identification techniques (Arnau-Gonzalez et al., 2021a; Kumar et al., 2021) use EEG features to identify certain persons among a large number of samples, which should be robust across tasks and sessions. In addition, SEED IV (Zheng et al., 2019) has four sessions of EEG data for EEG-based emotion studies, Langer et al (Langer et al., 2017) presented a dataset combining electrophysiology and eye-tracking intended as a resource for the investigation of information processing in the developing brain, and Kappenman et al (Kappenman et al., 2021) developed ERP Core with six tasks for ERP teaching studies.

**Table 1.** Open access EEG databases with characteristics of multi-session, multi-subject, or multi-task.

| Database | # Channels | Equipment | # Sessions | # Subjects | # Tasks | Main Usages |
| --- | --- | --- | --- | --- | --- | --- |
| Cho et al., 2017 | 64 | ActiveTwo / Biosemi | 1 | 52 | Rest, MI | Motor-BCI |
| Goldberger et al., 2000 | 64 | BCI2000 | 1 | 109 | Rest, MI, ME | Motor-BCI |
| BCI Competition IV-2a, 2008 | 22 | Unknown | 2 | 9 | MI | Motor-BCI |
| SEED IV 2019 | 62 | EPOC / Emotiv | 4 | 15 | Watch movie clips | Emotion Recognition |
| Korczowski et al., 2019 | 32 | g.USBamp / Gtec | 3 | 50 | P300 | P300-BCI |
| Lee et al., 2019 | 62 | BrainAmp / BrainProduct | 2 | 54 | Rest, MI, P300, SSVEP | BCI inefficiency |
| Jeong et al., 2020 | 60 | actiCHamp / BrainProduct | 3 | 25 | 11 different upper-extremity movement tasks | Motor-BCI |
| ERP Core 2021 | 30 | ActiveTwo / Biosemi | 1 | 40 | Rest, N170, MMN, N2pc, N400, P3, LRP and ERN | ERP studies |
| Langer et al., 2017 | 128 | Geodesic Hydrocel system | 2 | 126 | Passive: Rest, P-SS, NS, Active: CCD, SL, A-SS | Information processing in the developing brain |
| Gaspar et al., 2011 | 128 | ActiveTwo / Biosemi | 10 | 5 | ERPs (faces and noises) | ERP reliability |
| Kumar et al., 2021 | 128 | GES400 / EGI | 2-5 | 30 | OB, FUW, IBA, MMI, PA, SSVEP, PAV | personal identification |
| BED 2021 | 14 | EPOC / Emotiv | 3 | 21 | Rest, AS, VC<sub>x</sub> and VF<sub>x</sub> | personal identification |
| M<sup>3</sup>CV 2021 [this work] | 64 | BrainAmp / BrainProduct | 2 | 106 (95 for 2 sessions) | Rest, VEP, AEP, SEP, SSVEP, SSAEP, SSSEP, P300, ME, and SSVEP-SA | Intra- and inter-subject variability and cross-task analysis |

Abbreviations: Rest – Resting-state, MI – Motor Imagery, ME – Motor Execution, LRP – Lateralized Readiness Potential, ERN – Error-related Negativity, P-SS - Passive Surround Suppression, NS - Natural Stimuli, CCD - Contrast Change Detection, SL - Sequence Learning,

EEG-based biometrics is more difficult to be stolen, or forged and must be acquired by live detection. It can provide a more secure biometric method for identity recognition compared with existing biometrics, such as iris, fingerprint, and face. The research on EEG-based biometrics has received increased attention in recent years (Wang et al., 2020; Chan et al., 2018; Debie et al., 2021; Jin et al., 2021; Marcel and Millan, 2007). However, there are many challenges such as temporal permanence and robustness to mental state changes in an EEG-based biometric system (Cahn and Polich, 2006; La Rocca et al., 2013; Maiorana, 2021a; Maiorana and Campisi, 2018). At the same time, there is no open EEG-based biometric competition. The lack of a unified test benchmark and platform hinders the development of this field. To this end, we decided to open the M^3^CV dataset to launch an EEG-based biometric competition.

The remainder of this paper is organized as follows. Session II introduces the M^3^CV dataset. The details of the EEG-based biometrics competition are presented in Section III. Results are provided in Section IV. The discussion and conclusion are given in Section V.

## 2. M^3^CV Dataset

### 2.1 Subjects

A total of 106 healthy subjects from Shenzhen University participated in this experiment. Of these, 95 subjects (Age: 21.3 ± 2.2 years; Gender:73 males) participated in two sessions of the experiment, which were scheduled on different days, the between-session time is in the range of 6 days to 139 days, with a mean of 20 days. All subjects had the normal hearing, normal or corrected-to-normal vision, and no history of neurological injury or disease (as indicated by a self-report). During the experiment, the subjects were seated in comfortable chairs and kept about one meter from the screen.

Ethical approval of the study was obtained from the Medical Ethics Committee, Health Science Center, Shenzhen University (No. 2019053). All subjects were informed of the experimental procedure, and they signed informed consent documents before the experiment, in which they agreed to make their data open to access for research aim on the premise of concealing their personal information.

### 2.2 Experimental paradigm

Fig. 1 shows that the whole experiment was arranged in two sessions on separate days. The entire experiment with 6 paradigms, 15 runs, and 14 tasks was completed within 2 hours (around 50 mins of recording time and 70 mins for experiment preparation and rest between the consecutive runs).

**Fig. 1.**
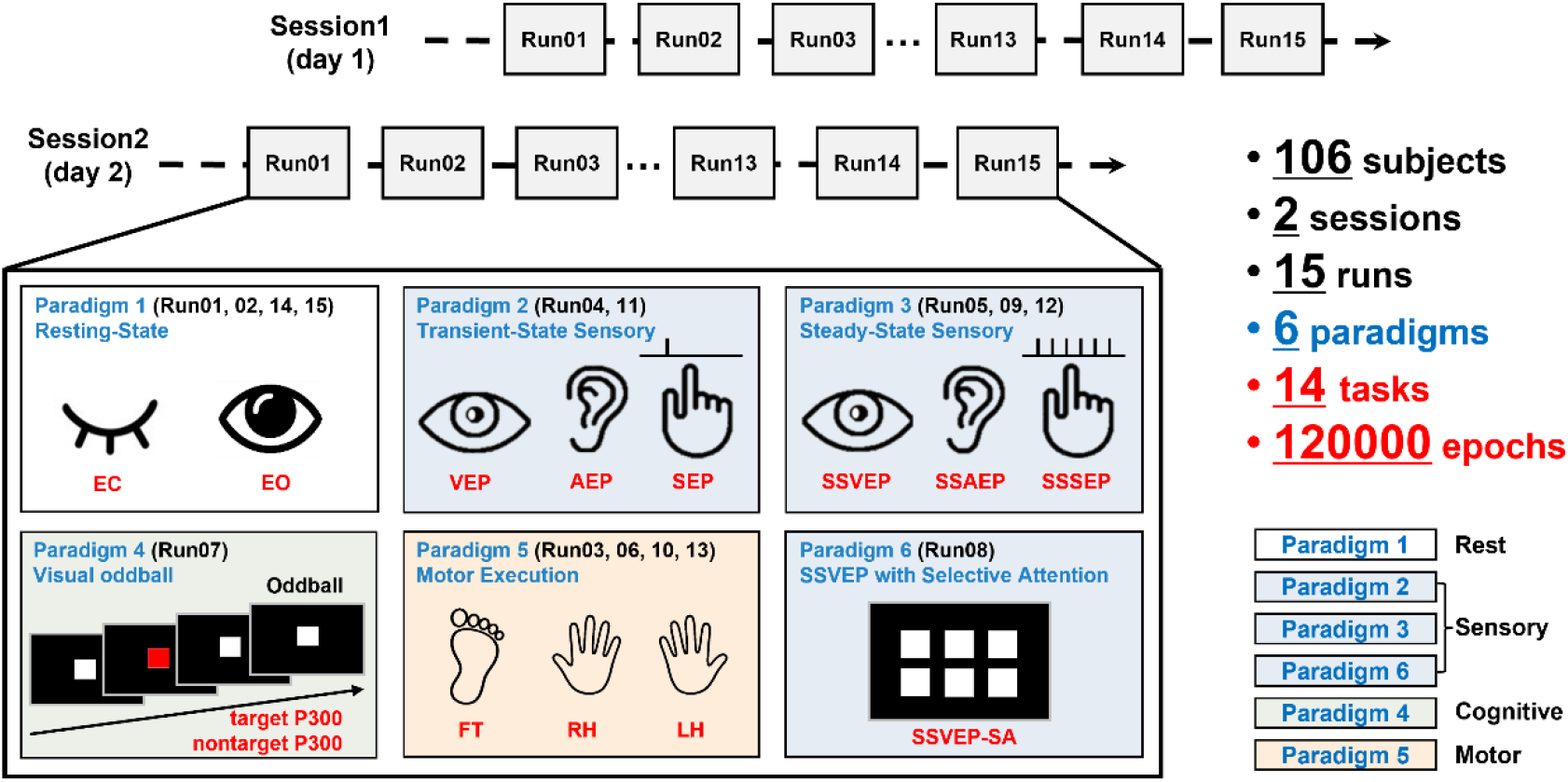
The experiment paradigm of the M^3^CV database.

The description of the 6 experimental paradigms is given below

1. **Resting-state**: Runs 01 and 14 were resting-state EEG signals with eyes-closed (EC), while Runs 02 and 15 were resting-state EEG signals with eyes-open (EO). Each lasting for 1 min. Instead of the fixation cross on the screen, the subject was asked to keep their eyes fixed on the LED (staying off) during the resting state with their eyes open. In the eyes-open run, the subjects were required to keep their eyes fixed in front and blink as little as possible.
2. **Transient-state sensory**: EEG elicited by visual, auditory, and somatosensory stimuli were recorded in Runs 04 and 11. For each run, 30 trials of VEP, AEP, and SEP were arranged in random order. Each stimulus lasted 50 milliseconds, and the inter-stimulus interval (ISI) was set at 2–4 s. On average, each run lasted 4.5 mins.
3. **Steady-state sensory**: A train of visual, auditory, and somatosensory stimuli were released in Runs 05, 09, and 12 respectively. Considering the different frequency responses of different modality stimuli, and the higher signal-to-noise ratio of SSVEP compared to SSAEP and SSSEP, the stimulation frequencies and recording times were different. They were 10 Hz (Herrmann, 2001) and 1 min for SSVEP in Run 05, 45.38 Hz (Galambos et al., 1981) and 2 mins for SSAEP in Run 09, and 22.04 Hz (Snyder, 1992) and 2 mins for SSAEP in Run 12.
4. **Visual Oddball**: A visual oddball experiment was arranged in Run 07 with the red square as the target stimuli and a white square as the nontarget stimuli on the screen. Each square lasted 80 ms with the ISI 200 ms. Six hundred trials of stimuli were arranged in 2 mins, in which target stimuli had a possibility of 5% (30 trials of target stimulation and 570 trials of nontarget stimulation). The subjects were asked to count the number of red squares and report after the run so that their attention would remain on the screen.
5. **Motor Execution**: Executed movement was performed in Runs 03, 06, 10, and 13. During these runs, the subjects were instructed to respond to a visual cue by gripping their left hand (LH) or right hand (RH), or by lifting their right ankle (FT) for a duration of 3 s, i.e., until the cue offset. No feedback was provided during the online recording. To ensure their motor areas were being fully activated, the subjects were required to perform the real executed movements of FT, RH, and LH at a rate of twice per second or faster, at approximately 80% of their maximum voluntary contraction, while keeping their upper body still. There was no external tool, like a metronome or hint on the screen, to remind the subject, because it may have produced unnecessary evoked potential as an external stimulus. During the experiment, the experiment monitored the movements of the subjects. If the subjects did not move fast enough or were found that their body moved simultaneously, the experimenter would abandon the recording of the current run, correct the subject’s movement and let them practice several more trials. The experimenters continuously monitored whether the movements of the subjects met these standards and corrected them when necessary. To minimize the intra- and inter-subject variability, we tried to ensure the consistency of the movements for all subjects. Hence, the executed movement was used instead of motor imagery in classical BCI experiments, and no feedback or training was given to the subjects.
6. **SSVEP with selective attention**: Six white squares flashed simultaneously at different frequencies in Run 08. First, the target square turned red for 200 ms. After 500 ms, all squares with the frequency of 7 Hz, 8 Hz, 9 Hz, 11 Hz, 13 Hz, and 15 Hz began to flash and lasted 10 s. The subject was asked to focus on the target square during the 10 s by covert visual attention with a fixation on the middle of the screen. In total, there were 12 trials arranged in Run 08. For each session of the experiment, they were instructed to perform 6 types of experimental paradigms, which were arranged in 15 runs, in which 14 tasks with rest, sensory, cognitive, and motor related EEG signals were listed in Table 2.

**Table 2.**
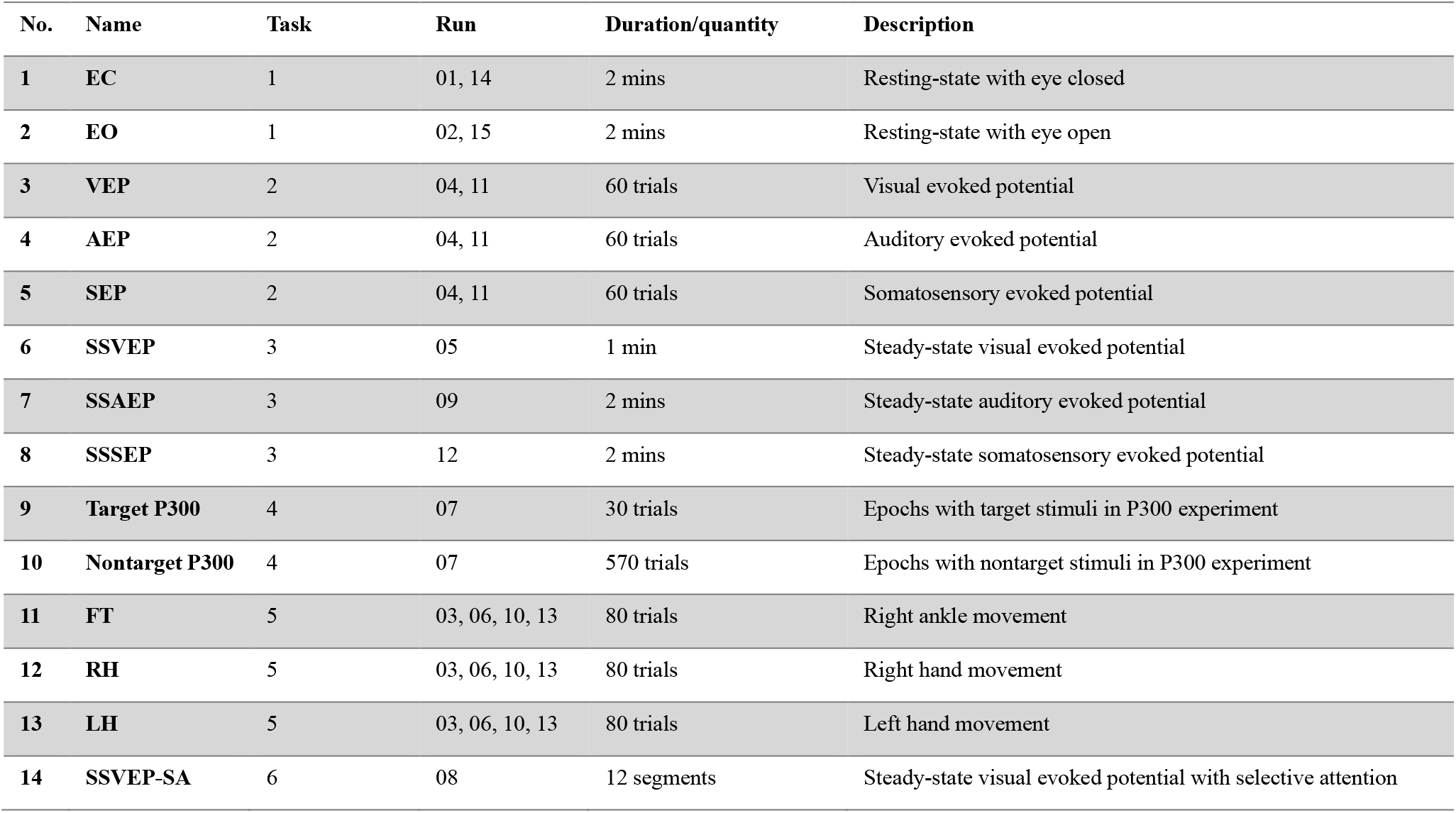
Description of 14 types of EEG tasks recorded in the M^3^CV database.

| No. | Name | Task | Run | Duration/quantity | Description |
| --- | --- | --- | --- | --- | --- |
| 1 | EC | 1 | 01, 14 | 2 mins | Resting-state with eye closed |
| 2 | EO | 1 | 02, 15 | 2 mins | Resting-state with eye open |
| 3 | VEP | 2 | 04, 11 | 60 trials | Visual evoked potential |
| 4 | AEP | 2 | 04, 11 | 60 trials | Auditory evoked potential |
| 5 | SEP | 2 | 04, 11 | 60 trials | Somatosensory evoked potential |
| 6 | SSVEP | 3 | 05 | 1 min | Steady-state visual evoked potential |
| 7 | SSAEP | 3 | 09 | 2 mins | Steady-state auditory evoked potential |
| 8 | SSSEP | 3 | 12 | 2 mins | Steady-state somatosensory evoked potential |
| 9 | Target P300 | 4 | 07 | 30 trials | Epochs with target stimuli in P300 experiment |
| 10 | Nontarget P300 | 4 | 07 | 570 trials | Epochs with nontarget stimuli in P300 experiment |
| 11 | FT | 5 | 03, 06, 10, 13 | 80 trials | Right ankle movement |
| 12 | RH | 5 | 03, 06, 10, 13 | 80 trials | Right hand movement |
| 13 | LH | 5 | 03, 06, 10, 13 | 80 trials | Left hand movement |
| 14 | SSVEP-SA | 6 | 08 | 12 segments | Steady-state visual evoked potential with selective attention |

### 2.3 Experimental platform

The continuous EEG signals were recorded using an EEG amplifier (BrainAmp, Brain Products GmbH, Germany) and multichannel EEG caps (64 Channel, Easycap). The signals were recorded at a sampling rate of 1000 Hz by 64 electrodes, placed in the standard 10–20 positions. The electrodes FCz and AFz served as reference and ground, respectively. Before data acquisition, the contact impedance between the EEG electrodes and the cortex was calibrated to be lower than 20 kΩ to ensure the quality of EEG signals during the experiments.

An Arduino Uno platform was programmed to release the visual, auditory, and somatosensory stimuli in tasks 2 and 3, which communicated with the Matlab program (The MathWorks Inc., Natick, USA) on a PC through a serial port.

- Visual stimuli for VEP and SSVEP were delivered by a 3W light-emitting diode (LED) with a 2cm diameter circular light shield, which was placed in the center of the visual field of view 45cm away from the subjects’ eyes. The mean LED intensity was 1074 Lux as measured by a light meter (TES-1332A, TES).
- Auditory stimuli for AEP and SSAEP were presented via a Nokia WH-102 headphone. 1000Hz pure tone was applied for the stimuli. The intensity was set at a comfortable level (75 dB SPL on average) for all subjects as measured by a digital sound level meter (Victor 824, Double King Industrial Holdings Co., Ltd. Shenzhen, China).
- Somatosensory stimuli for SEP and SSSEP were generated by a 1027 disk vibration motor. Since there was no effective tool to measure the output intensity of the vibrator directly, we have to report the detail of the product parameters with the rated power of 3W, efficiency of 80%, and dimensions of 10mm*2.7mm).

For the other tasks, A 24.5-inch screen (1920*1080) with a 240-Hz refreshing rate (Alienware AW2518H, Miami, USA) was used to present the visual stimuli or cues by programmed using Psychtoolbox-3 (http://psychtoolbox.org/) in Matlab. The red and white squares were delivered in a sequence at the center of the screen with a black background for visual oddball. Three white squares were also used in the motor execution paradigm to indicate the movement. Six white squares flashed simultaneously at different frequencies to deliver the stimulus in SSVEP with selective attention paradigm. The size of the square was set to be 300*300 pixels in these paradigms.

### 2.4 EEG signal pre-processing

The data pre-processing pipeline on the M^3^CV database is illustrated in Table 3. The raw EEG signals were recorded with BrainVision core data format (each recording consisting of a .vhdr, .vmrk, .eeg file triplet), which were managed with Brain Imaging Data Structure (BIDS) (Gorgolewski et al., 2016; Pernet et al., 2019). For each recording, the bad channels were interpolated first. Channel FCz (the reference) was added back, and channel IO was removed. Then all signals were filtered by a 0.01–200-Hz band-pass filter and a 50-Hz notch filter. A 2-order Butterworth zero-phase filter was applied in the above two steps of filtering. After re-referencing to TP9/TP10, artifacts produced by eye blinks or eye movements were identified and removed manually by Independent Component Analysis (ICA) (Huang et al., 2020).

**Table 3.** Pipeline for EEG pre-processing.

| Software | Matlab 2018b & Letswave7 (Letswave.cn) |
| --- | --- |
| Band-pass filtering | Butterworth filter, 0.01-200 Hz, 4<sup>th</sup> order, 24dB/octave, zero-phase |
| Notch filtering | Butterworth filter, 49-51 Hz, 4<sup>th</sup> order, 24dB/octave, zero-phase |
| **Channel interpolation** | Bad channels were identified manually and interpolated with the mean value of the three surrounding channels |
| **Re-reference** | Re-reference to the mean value of TP9 and TP10 |
| **Artifacts removal by ICA** | Eye movement related ICA components were identified by visual inspection of their scalp topographies, time courses, and spectra. |

During the signal preprocessing, we interpolate bad channels for 22 subjects and one subject was removed due to strong 10 Hz artefacts among the 95 subjects who complete the two sessions. No bad epoch has been removed. For Motor execution, preventing the interference of EMG artifacts caused by actual movement was crucial during our data collection, especially in the movement paradigm. Firstly, we asked the subject to keep their torso still in both foot and hand movement. That is also the reason why we set right foot movement instead of two-foot movement in the experiment design. Secondly, the experimenters continuously monitored the subjects’ movement and online EEG display to ensure the quality of data collection. Thirdly, since the amplitude of EMG single is much higher than the EEG signal, the contamination of EMG artifacts is highly visible. Also, we did not see strong EMG artifacts during the signal preprocessing. Furthermore, no strong EMG artifact is observed in the time-frequency analysis. The main EEG responses came from C3, C4, and Cz at *μ* and *β* band, The *γ* band response was much smaller, which was the main frequency band for EMG artifacts.

After experiment, in the preprocessing process we interpolate bad channels for 22 subjects and one subject was removed due to strong 10 Hz artefacts among the 95 subjects who complete the two sessions.

### 2.5 Basic EEG signal visualization

To show the basic information of EEG signal in M^3^CV, we perform time domain analysis on ERP signal, Frequency domain analysis on resting-state EEG and steady-state evoked potentials, and time-frequency domain analysis on motor-related signals.

1. **Time domain analysis**. The ERPs of VEP, SEP, AEP, and target and non-target P300 were analyzed in the time domain. A Butterworth bandpass filter with 0.5 – 30 Hz was applied to the preprocessed EEG signal. After segmentation and averaging, baseline correction was performed from −0.5 to 0 s to obtain the ERP for each subject.
2. **Frequency domain analysis**. For the task of resting-state with eye closed and eye open, the EEG signals were firstly re-referenced to a common average reference after preprocessing. Welch’s method with a 2-s window and 50% overlap was applied. After transforming the total power level into decibels and averaging, the frequency domain responses for resting-state EEG with eye closed and eye open were obtained for each subject. The processing pipeline for EEG signals from steady-state sensory and SSVEP with selective attention tasks was similar. But no re-reference was performed, and TP9/TP10 was still used as the reference. FFT was applied instead of Welch’s method to obtain a sharp frequency response for SSVEP, SSAEP, SSSEP, and six classes of SSVEP-SA signals.
3. **Time-frequency domain analysis**. For the motor execution tasks, since the EEG responses were complex in different time intervals and frequency bands, a continuous wavelet transform was applied for the time-frequency domain transform, in which the complex Morlet wavelet was used with the central frequency 1.5 Hz and bandwidth 1 Hz. Considering the computation complexity, the EEG signal was downsampled to 200 Hz to reduce the time for computation. After the continuous wavelet transform, the time-frequency response was further downsampled to 50 Hz to reduce the storage space on the hard disk and computer memory.

## 3. EEG-based biometrics

### 3.1 Literature review

Several studies have reported that EEG-based identification and verification systems are capable of high recognition accuracy (Kong et al., 2019; Koike-Akino et al., 2016). However, many of them have not been evaluated under more rigorous conditions, such as testing with large population size, across sessions, and across tasks, which were detailed as follows:

- **Uniqueness:** Although many studies have revealed the uniqueness of EEG signals and their ability to discriminate between subjects, the tested databases normally have a small population size of fewer than 50 subjects (Delpozo-Banos et al., 2018; Arnau-Gonzalez et al., 2021b; Chen et al., 2016). For employing EEG as a biometric trait, it must be tested on a larger database.
- **Permanence:** EEG signals could be different in days even for the same task, which poses a major challenge to decoding subject identity (La Rocca et al., 2013; Maiorana et al., 2016; Maiorana and Campisi, 2018). Further, in single-session settings, machine learning algorithms may recognize the special electrode situation and preprocessing parameters rather than the EEG-based personal trait.
- **Robustness:** For the same session, EEG signals could vary significantly under different tasks or experimental conditions. A good biometric system should be robust to the above conditions (Wang et al., 2019; Maiorana, 2021a; Kumar et al., 2021; Kong et al., 2018), otherwise, an enrolled subject may be rejected or misidentified by the system when they are under different mental states.

According to the three points mentioned above, we did a short review of the existing studies on multi-session and multi-task EEG-based biometrics as shown in Table 4. Compared with our proposed database, the population size of these studies is quite small, especially for (La Rocca et al., 2013) and (Zeynali and Seyedarabi, 2019). As for feature extraction, hand-crafted features, such as AR, MFCC, and PSD, are still the dominant choice while only Maiorana (Maiorana, 2020) and Kumar et al (Kumar et al., 2021) applied deep learning methods. In terms of performance metrics, the recognition accuracy and the equal error rate were the most used metric for identification tasks and verification tasks respectively(Arnau-Gonzalez et al., 2021a; Kumar et al., 2021). It should be noted that these studies were performed on multi-session and multi-task EEG databases, but only Kumar et al (Kumar et al., 2021) applied cross-session and cross-task evaluation to their proposed algorithm.

**Table 4.**
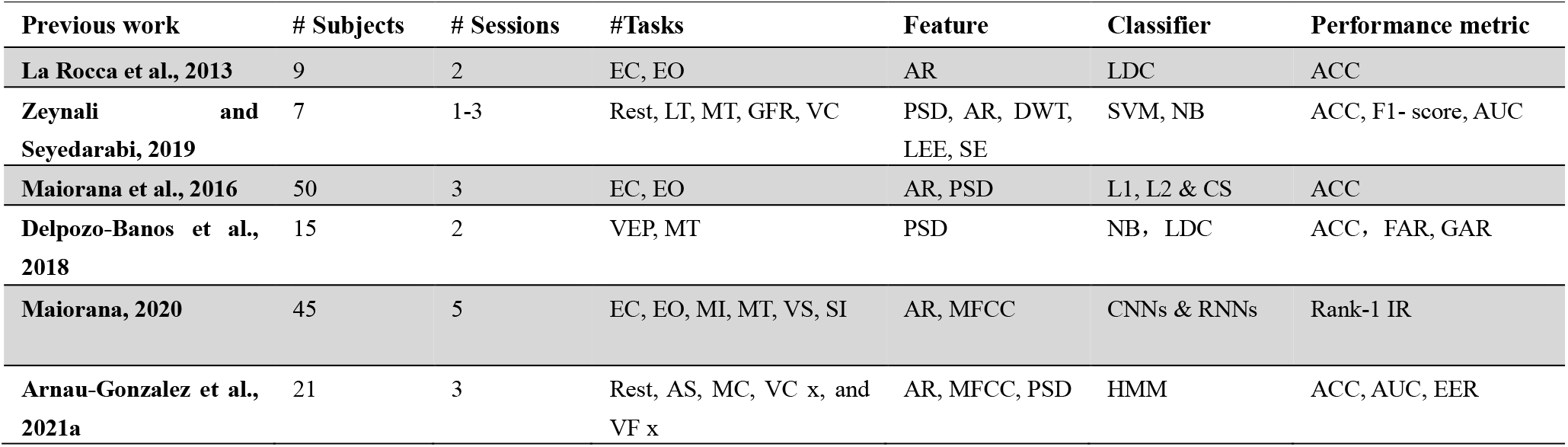
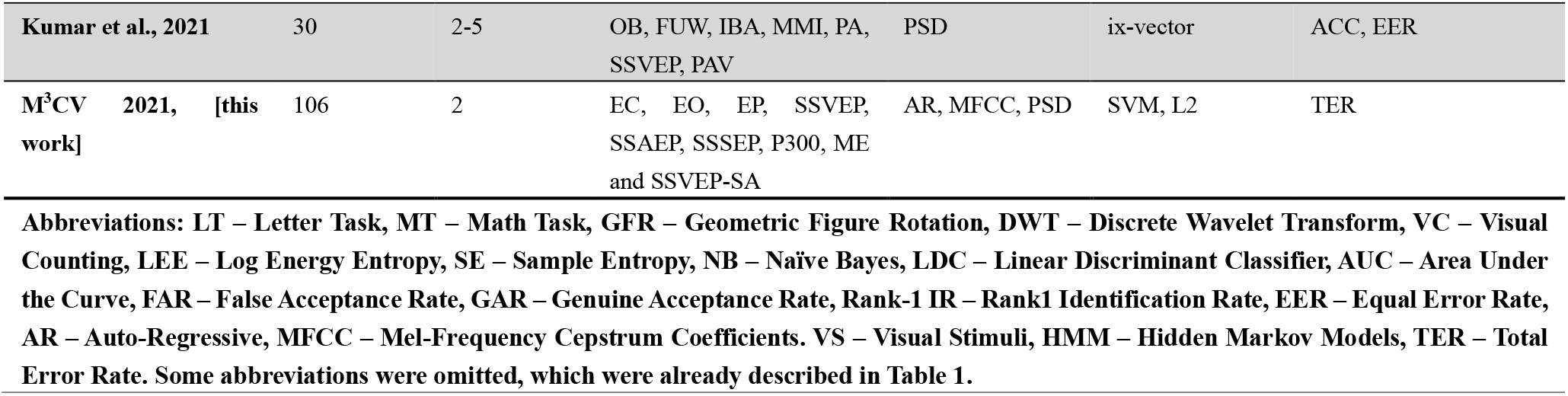
Literature review on multi-session and multi-task EEG-based biometrics study

### 3.2 Learning Task

Based on the M^3^CV database, the EEG-based biometric competition was launched by focusing on the problems of personal identification and verification (Fig. 2). The contestant needs to train the classification model for the biometric system with the enrollment dataset. After the model has been trained, the contestant would be asked to complete the following two classification tasks:

**Fig. 2.**
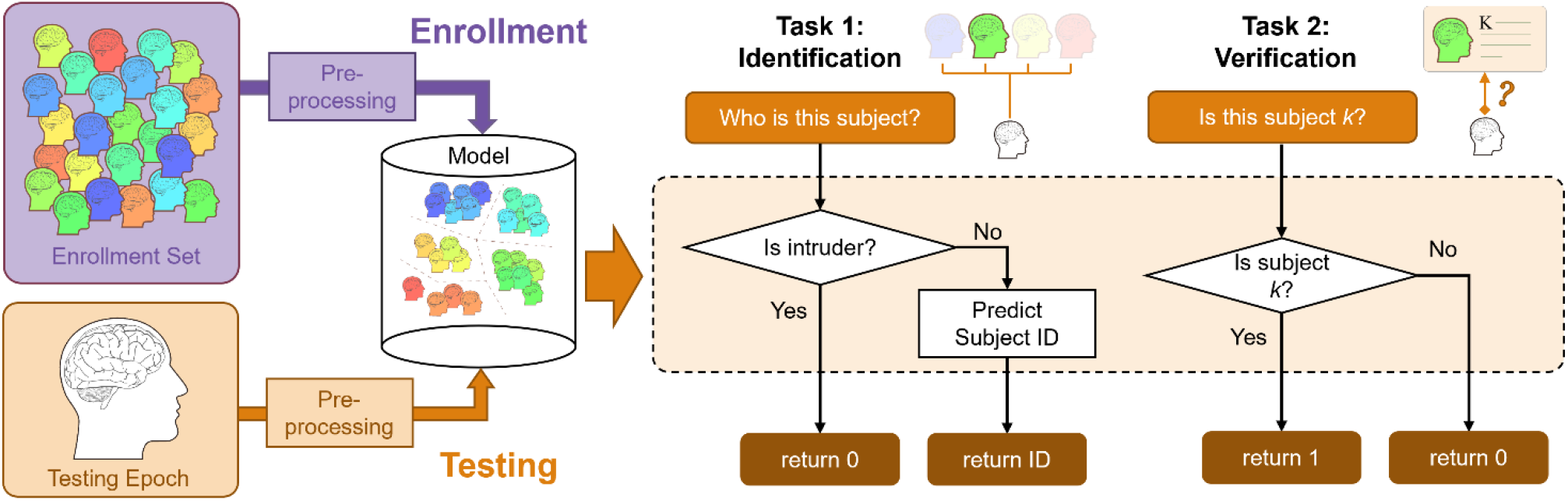
The architecture of the EEG-based biometric system with the learning task of personal identification and verification.

**Fig. 3.**
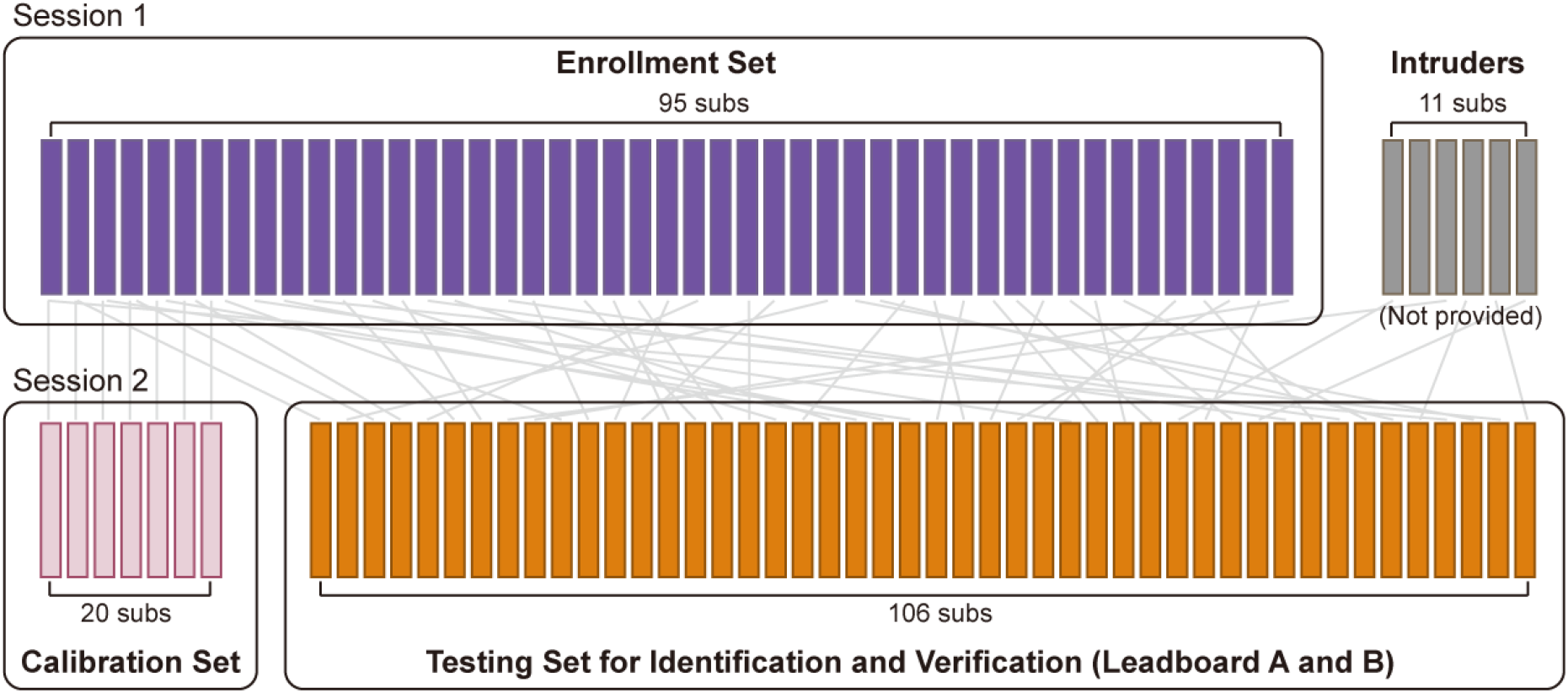
The segmentation of M3CV for online competition. The whole set was divided into three parts: Enrollment Set, Calibration Set, and Testing set (containing 11 intruders).

- **Identification:** Determine whether a given EEG signal comes from the enrollment set, and further determine the subject ID if the epoch is not judged as from an intruder. The potential applications of identification systems include video surveillance and information retrieval (e.g., identifying criminals from police databases).
- **Verification:** Determine whether a given EEG signal comes from a certain subject. The potential applications of the verification system are to access control systems (airport checking, monitoring, computer or mobile devices log-in).

### 3.3 Baseline model

#### 3.3.1. Feature extraction

According to the literature review, three types of commonly used features were selected for EEG-based biometrics based on the EEG signal pre-processing:

1) Power Spectral Density (PSD): To estimate the power spectrum of each channel for each epoch, we used the Welch periodogram algorithm. Specifically, we divided the whole 4s epoch into 1s segments with 0.5s overlap, then we averaged the FFT of each segment using a hamming window. The PSD spectrum from 2 to 45 Hz was evenly divided into 12 frequency bands, then the band power of these frequency bands was extracted.

2) Mel Frequency Cepstral Coefficients (MFCCs): MFCCs are one of the most common features of speech recognition, and have been recently applied to the biometric analysis of EEG data (Arnau-Gonzalez et al., 2021b; Maiorana, 2020). MFCCs are extracted from the mel-filtered spectrum amplitudes *s*_*m*_ using log compression and discrete cosine transform, defined as

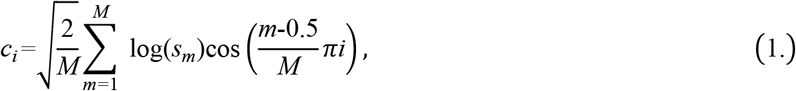

where L is the number of preserved cepstral coefficient, *c*_*i*_ is the *i*-th cepstral coefficient, with *i* = 1,2,3, …, *L. M* is the number of triangular band-pass filters in a mel-scaled filter bank and *L* is the number of the first cepstral coefficients used to obtain signal representation. In this study, *M* = 18 and *L* = 12 were employed.

3) Autoregressive Coefficient (AR): AR features are commonly used for EEG-based identification and verification (Maiorana and Campisi, 2018). In this study, AR features were extracted from each epoch from a 12-th order autoregressive model created by solving the Yule-Walker equations. Thus 12 AR coefficients were extracted for each EEG channel.

#### 3.3.2 Classification

In the biometric system, the verification task is a one-to-one classification problem and the identification task is a one-to-N classification problem. With the existence of the intruders, identification becomes a one-to-(N+1) classification problem. Considering the extensibility of the enrollment set, using N one-to-one weak classifiers from each subject to compose a one-to-N classifier is a practical way for the biometric system. The method can effectively avoid the retraining of the whole classifier after the enrolling of new subjects.

In this study, one-to-one classifiers, including the similarity-based method based on Euclidean distances (L2) and the One-class Support Vector Machine (SVM), were applied as the baseline classifiers for both verification and identification tasks in the subsequent analysis for simplicity. More specifically, after extracting features, an L2-based or SVM classification model was trained by epochs from each subject in the enrollment set.

In the verification task, the corresponding classifier would confirm the subject ID claimed by test epochs when the acquired score was greater than the rejection threshold *δ*. In an offline system, this threshold was varied to obtain the equal error rate (*V*_*EER*_), which was a commonly used threshold-independent measure in biometric verification tasks (Bidgoly et al., 2022), defined as the intersection point of false acceptance rate (*V*_*FAR*_) curve and false rejection rate (*V*_*FRR*_) curve. In an online system, the threshold has to be pre-determined, which was normally decided by obtaining EER on the training set (Marcel and Millan, 2007; Kang et al., 2018).

In the identification task, the intruder test was performed firstly, in which the test epoch was fed into each subject’s classification model to generate a score, when the maximal score obtained from all subjects’ models was below the threshold, the test epoch was judged as an intruder. For the L2-based model, the rule of judging the test epoch as an intruder was defined by Eq. (2)

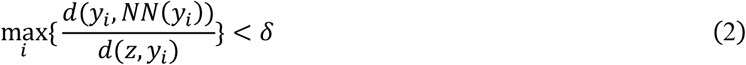

where *N* is the number of subjects in the enrolled set, *z* is the test epoch, *y*_*i*_ is the nearest epoch of *z* from all epochs of *i*-th subject with *i* = 1,2,3, …, *N, NN*(*y*_*i*_) is the nearest epoch of *y*_*i*_ from all epochs of *i*-th subject, *δ* is the rejection threshold determined by *V*_*EER*_. The subject ID would be assigned based on the maximum score from all subjects if the test epoch *z* passed the intruder detection. For an SVM-based classifier, the likelihood is applied instead of the distance in the L2-based classifier.

### 3.4 Online competition

1. **Dataset Composition:** The whole set for this competition was divided into three parts: Enrollment Set, Calibration Set, and Testing set (containing 11 intruders).
  - Enrollment Set: it provides the 1^st^ session of EEG from 95 subjects. The enrollment set is used for the contestants to train the biometric model.
  - Calibration set: the EEG data of the 2^nd^ session from 20 subjects is provided in the calibration set, which is used to help the contestants get familiar with the data format, refine effective feature extraction, and tune hyper-parameters of the machine learning model. The number of subjects in the calibration set is relatively small as compared with the enrollment set and testing set.
  - Testing set: it is used to evaluate the performance of the algorithm in the competition with the leaderboards A (public) and B (private), for which epochs belonging to leaderboard A or B are hidden. The testing epochs come from the 2nd session of 106 subjects, among which 95 subjects also exist in the enrollment set, and the other 11 subjects are treated as an intruder. And it should be noted that several epochs in the testing set come from the same subjects as that in the calibration set, but there is no overlap of the epochs.
2. **Organization of the competition:** We would launch the challenge on June 15^th^, 2022, and close it on Aug 25^th^, 2022. We would award money prizes to the top 9 contestants and certificates of honor for the 20 best contestants. Challengers were ranked based on the score of recognition accuracy of their submission computed on the private leaderboard (Leaderboard B). We framed the challenge problem by providing: (i) an open accessed dataset, (ii) a standard way to assess the submission, and (iii) a starting kit. For this purpose, we used the PaddlePaddle AI Studio platform. During the challenge, contestants submitted their solutions on the PaddlePaddle AI Studio website (https://aistudio.baidu.com/aistudio/competition/detail/315/0/introduction), and we provided to the contestants the score computed on the public leaderboard (Leaderboard A). At the end of the challenge, we asked each contestant to select a single submission with the code and description document. Contestants were ranked based on the score of these submissions computed on the private leaderboard (Leaderboard B).
3. **Competition platform:** PaddlePaddle’s AI Studio is a feature-rich, open-source, industry-level deep learning platform. With the full-process function and professional service support, the platform has held more than 150 AI competitions, attracted 500,000 elite developers from many countries around the world, and trained tens of thousands of top players. It is committed to creating the world’s leading AI competitions, gathering talents, and promoting the development of the intelligent economy.
4. **Data and code availability:** The public dataset with the code of the baseline methods and an example submission file are available at https://aistudio.baidu.com/aistudio/datasetdetail/151025/0.

### 3.5 Benchmarks

To make a comprehensive understanding of the challenges in the EEG-based biometrics competition, we have provided the benchmark test in three steps. 1) Offline evaluation was applied to investigate the influence of the increasing population size, cross-session, and cross-task tests on the performance of the biometric system. 2) Simulated online analysis was applied to evaluate the baseline methods by simulating the competition process with data from the enrollment set and calibration set. 3) Online submission shows an example submission with the baseline method with a score returned from competition platform.

#### 3.5.1 Offline evaluation

We performed three analyses to show the influence of the increasing population sizes, cross-session, and cross-task tests on the performance degradation of EEG-based biometrics. These analyses were done on the data from 20 subjects in the calibration set and enrollment set, whose subject ID of the second session was known. The reason of restrict our analyses to these 20 subjects is to avoid any information leakage about the testing set. The train/test pipeline and performance metric for these analyses were detailed as follows:

1. **Testing on increasing population sizes:** In this test, we increased the number of subjects from 2 to 20, with the increasing step of 2 subjects each time, to investigate the influence of increasing population sizes on the performance of EEG-based biometrics. To avoid bias, subjects were randomly selected 10 times for each number of subjects. For each task, the data from the first session was used to train, and the data from the second session was used to test. Since the aim of this test is to show the influence of increasing the number of subjects, the intruder is not considered.
2. **Testing across tasks:** In this test, EEG data of a particular task from one session (session 1/session 2) was used to train the model, and EEG data of another task from the other session (session 2/session 1) was used to test. And then the training set and the testing set would be exchanged. The population size was set to 20 in the test. Since AEP, SEP and VEP are in the same runs, we treated them as one condition/task, EP (i.e., Evoked Potential). Also, the term ME (Motor execution) was used to represent the LH, RH, and RF conditions.
3. **Testing across sessions:** In this test, to make a fair comparison of the performance between within-session and cross-session tests, we used the same training set for the within-session tests and cross-session tests, which is the first half of the epochs in the first session. The testing set for the within-session test is the second half of the epochs in the first session, while the testing set for the cross-session tests is all epochs from the second session. This procedure was done for each task separately. And the population size was set to 20 in the test.

For offline evaluation, the performance metrics consists of *I*_*ER*_ and *V*_*EER*_. For the identification task, the identification error rate *I*_*ER*_ is defined as

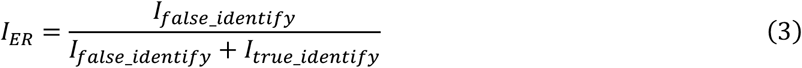

in which *I*_*false_identify*_ and *I*_*true_identify*_ are the number of epochs being wrongly and truly identified. For the verification task, since the subject ID of the testing set is known, the verification equal error rate *V*_*EER*_ was commonly used as a conventional performance metric in biometric verification tasks for offline analysis (Maiorana, 2021b). With the growth of the reject threshold in the classifier model, the false acceptance rate (*V*_*FAR*_) increases and false rejection rate (*V*_*FRR*_) decreases, and *V*_*EER*_ is the intersection point of the *V*_*FAR*_ and *V*_*FRR*_ curve. *V*_*FAR*_ and *V*_*FRR*_ are defined as

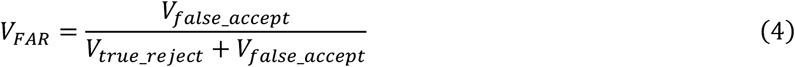

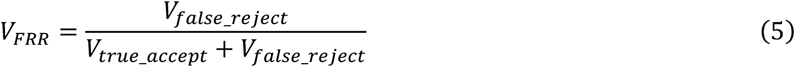

in which *V*_*false_reject*_ and *V*_*true_accept*_ are the number of epochs being wrongly rejected or truly accepted for whose true identity is the same as the claimed ID. Similarly, *V*_*false_accept*_ and *V*_*true_reject*_ are the number of epochs being wrongly accepted and truly rejected for whose true identity is not the same as the claimed ID.

#### 3.5.2 Simulated online analysis

During the offline evaluation, *V*_*EER*_ is commonly used as an important metric to evaluate the performance of the verification result. However, in the online application, the equation *V*_*FAR*_ = *V*_*FRR*_ on calibration set may not hold on to the testing set, the metric *V*_*EER*_ may not reflect the real performance of a biometric system. Furthermore, the time consumption would also be an important metric for a practical biometric system.

Therefore, beyond offline evaluation, we further applied simulated online analysis to evaluate the performance of different baseline methods of feature extraction and classification. Ten subjects in the calibration set and corresponding enrollment set were used to determine the optimal threshold, which was defined as the one that making *V*_*FAR*_ = *V*_*FRR*_. Another 10 subjects were used to evaluate the performance of the six baseline models for each combination of feature extraction (PSD, AR, MFCC) and classification (L2 and SVM) methods.

For simulated online analysis, the performance metrics of *I*_*ER*_, *V*_*ER*_, *V*_*FAR*_, *V*_*FRR*_, *T*_*train*_, and *T*_*test*_ were used in the evaluation, in which *T*_*train*_ and *T*_*test*_ are the time consumption for training and testing, and verification error rate *V*_*ER*_ is defined as

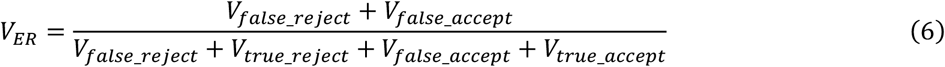

#### 3.5.3 Online submission

For online submission, an example code is provided with the methods of feature extraction (PSD, AR, MFCC) and classification (L2 and SVM), in which the optimal threshold was determined by making the *V*_*FAR*_ = *V*_*FRR*_ on the 20 subjects in the calibration set and enrollment set, whose subject ID of two sessions was known.

An example submission file was also provided with the method of PSD+L2. With the subject ID unknown in the testing set, only the score of the total recognition accuracy returned after submission can be used to evaluate the performance. The result can also be obtained if the contestant submits the same result file to the system. The code and the submission file are available at the link: https://aistudio.baidu.com/aistudio/datasetdetail/151025/0.

Hence, for online submission, the performance metric is the score as 1-total error rate (*TER*), which is computed by Eq. (7)

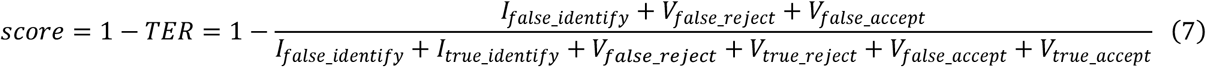

## 4. Results

### 4.1 Basic EEG results for visualization

The basic EEG results from the proposed M^3^CV database are shown in Fig. 4-6, which include the results from the time domain, frequency domain, and time-frequency domain.

**Fig. 4.**
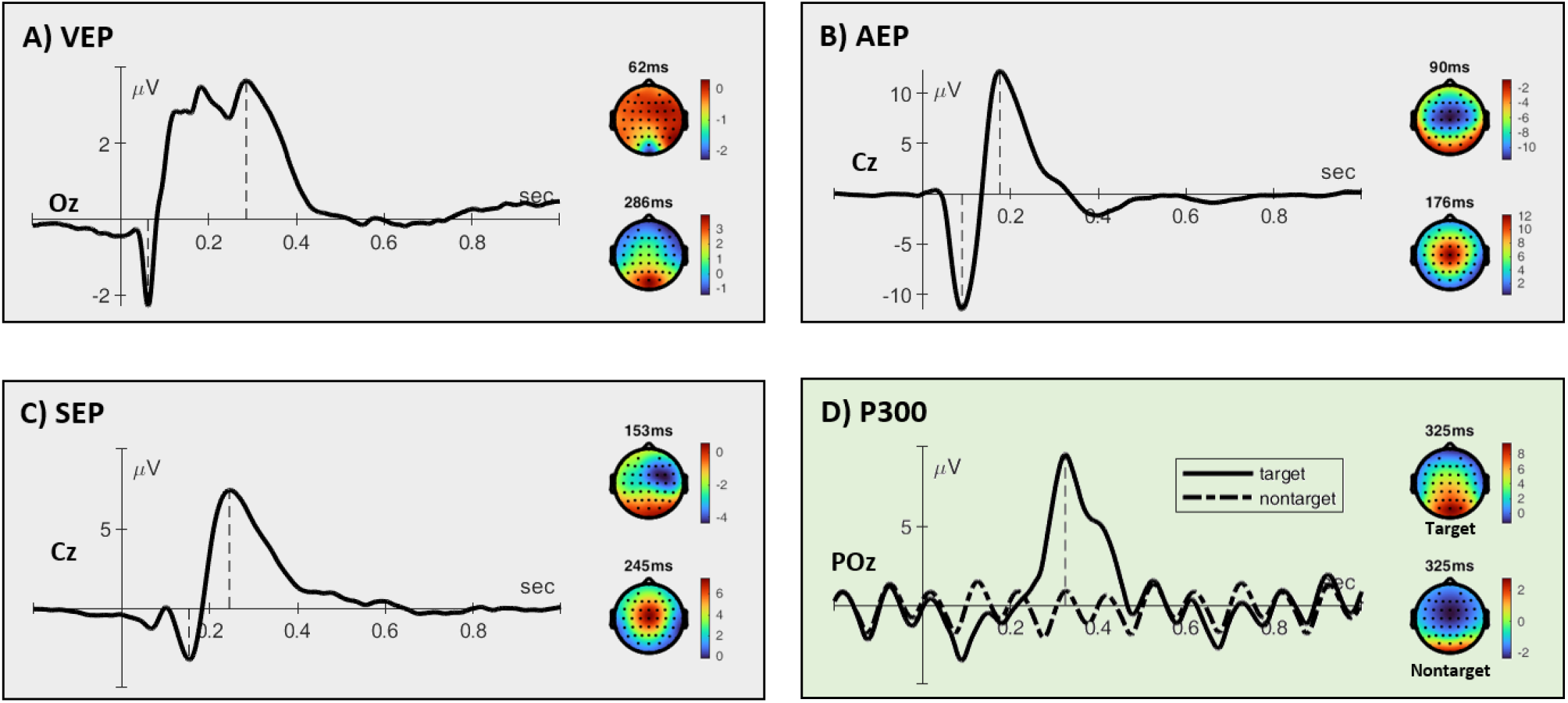
Time domain EEG results for A) VEP, B) AEP 2, C) SEP in paradigm 2 and D) P300 in paradigm 4.

For time-domain analysis. The grand-averaged waveforms of VEP, AEP, SEP, and target- and nontarget-P300 are shown in Fig. 4 (A–D), with the topographies at their corresponding peaks. The response of VEP mainly concentrated on the occipital area. AEP and SEP shared a similar N2/P2 waveform in the central area. A classical N75-P100-N135 complex is not seen here, which is typically observed from pattern-reversal VEP, for which high-contrast, black-and-white checkerboards are used as stimuli. As the vibrator was placed on the subject’s left hand, the N2 response of SEP showed a response from the contralateral primary sensory area. The topographies for the P2 components from AEP and SEP are quite similar. In the oddball experiment, the target P300 response was mainly concentrated on the channel POz at around 300 ms. By comparison, the nontarget response mainly came from the occipital area because the visual stimuli were used in the oddball experiment.

For frequency domain analysis of the resting-state EEG with EC and EO (Fig. 5A), the difference was concentrated on the occipital area. With the increase in frequency, the amplitude of the EEG response decreased at a rate of approximately 1/freq for both EC and EO. The main difference between EC and EO was the three peaks that appeared in sequence around 10 Hz, 20 Hz, and 30 Hz. For the first peak in the interval of 8-12 Hz, the main response was in the occipital area, the frequency power at Cz (0.45 dB) is larger than C3 (0.17 dB) and C4 (0.21 dB) in the condition of EO, while the frequency power at Cz (−1.05 dB) would be smaller than C3 (−0.88 dB) and C4 (−0.73 dB) in the condition of EC. The topographies for the second peaks in the interval of 18-22 Hz had a similar shape, while the difference mainly came from the magnitude. For SSEP analysis in Fig. 5B, the brain areas and frequency band of the induced SSEP were different for visual, auditory, and somatosensory stimuli, which were around 10 Hz at the occipital area for SSVEP, around 45 Hz at the frontal-central area for SSAEP, and around 22 Hz at the primary sensory area for SSSEP. Fig. 5C shows how attention modulated the foundation frequency (black triangles) and their harmonic (gray triangles) responses at different frequency points. With selective attention, the target frequency (downward-pointing triangles) responses were larger than those at other nontarget frequencies (upward-pointing triangles).

**Fig. 5.**
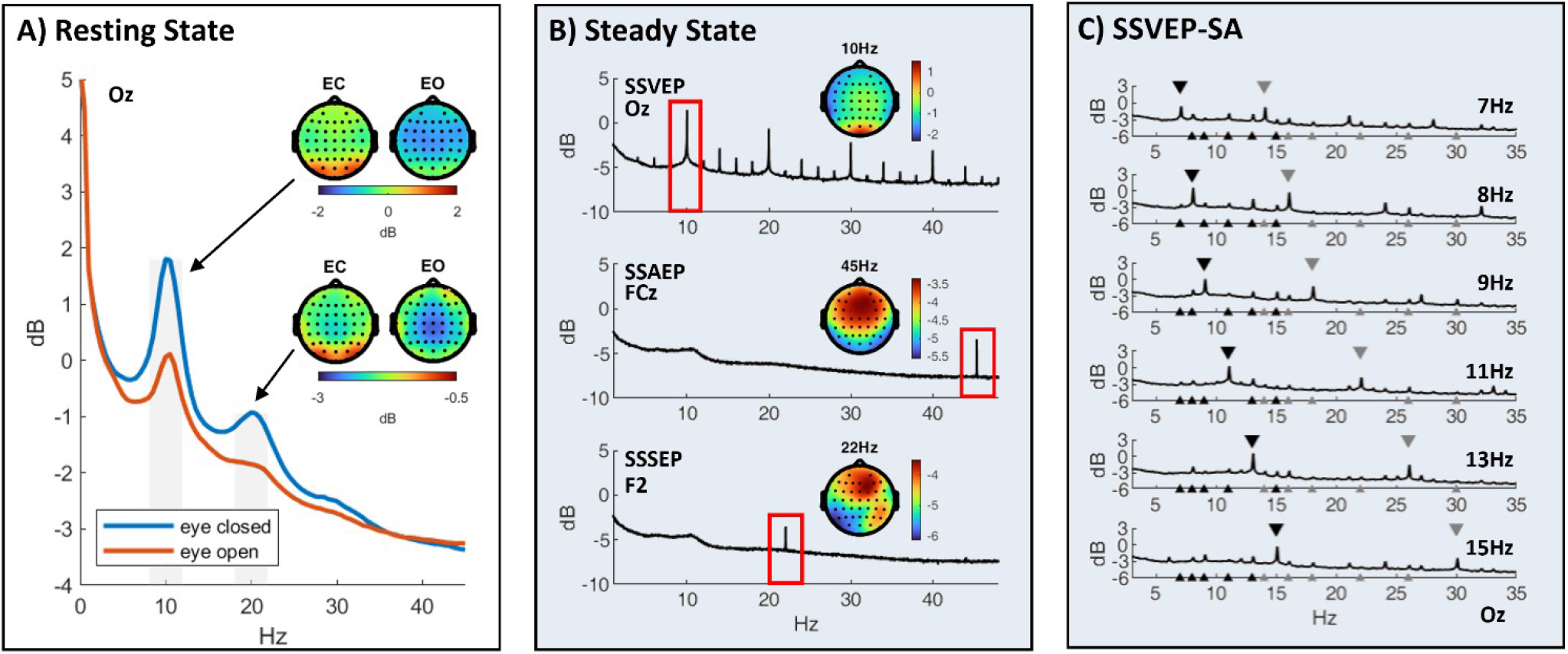
Frequency domain EEG results for A) Resting-state EEG with eye closed and eye open, B) Steady-state Evoked potential with SSVEP, SSAEP, and SSSEP, and C) SSVEP with selective attention.

Time-frequency analysis was performed for the motor task because foot and hand movements led to rich neural oscillation changes in different time-frequency-spatial regions. Fig. 6H is a comprehensive illustration of these changes caused by these three types of movements, which are FT, RH, and LH. Further, no strong EMG artifact is observed in the time-frequency analysis. The main EEG responses came from C3, C4, and Cz at *μ* and *β* band, The *γ* band response was much smaller, which was the main frequency band for EMG artifacts.

**Fig. 6.**
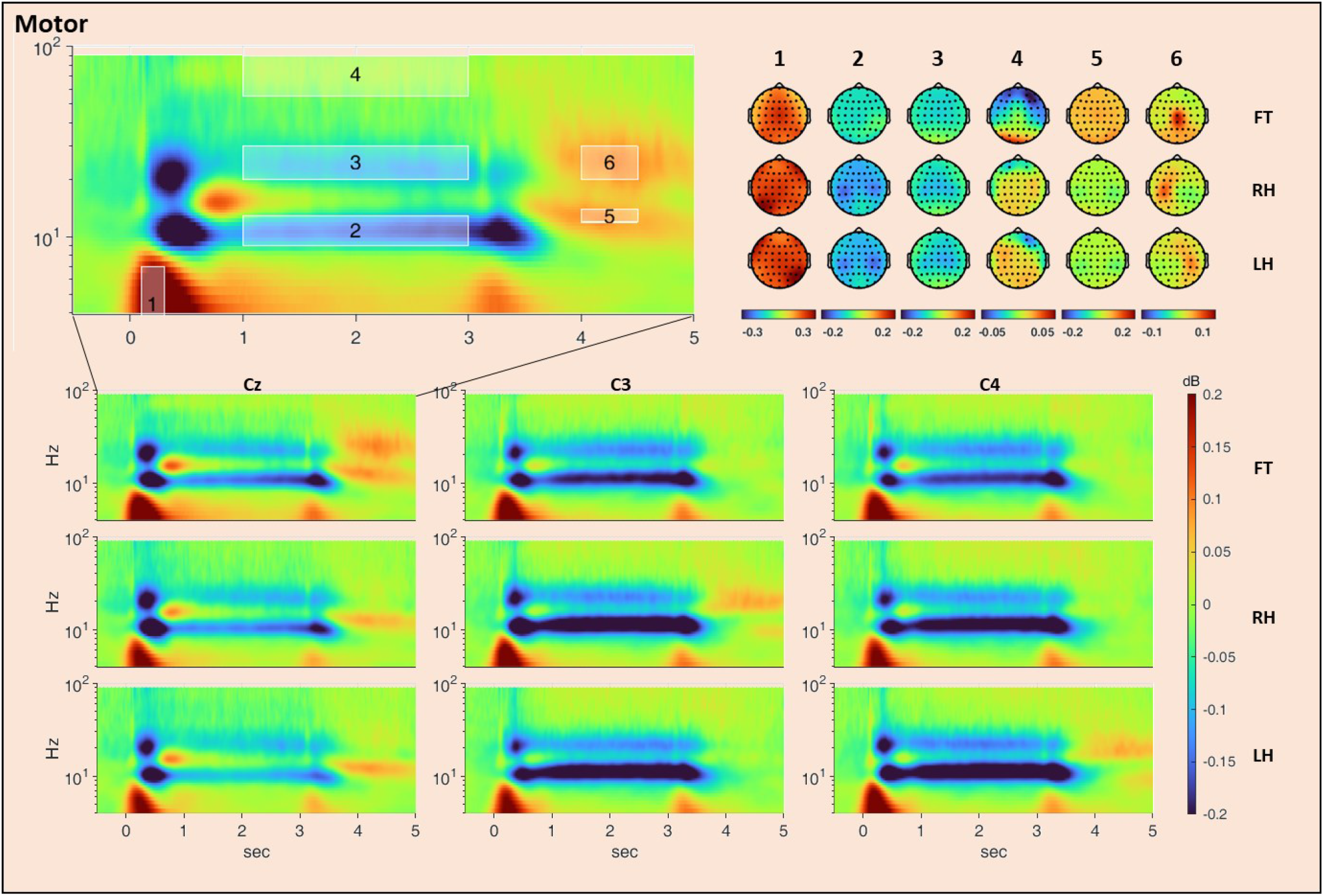
Time-frequency domain EEG results for motor execution task for FT (right angle movement), RH (angle hand), and LH (left hand movement) at channels Cz, C3, and C4. The topographies of the mean magnitude from the six typical regions are illustrated for three types of movements.

The corresponding topographies from six regions of interest (ROI) are detailed below.

- ROI #1 (from 0.1 to 0.3 s, 2 to 7 Hz): The response in this region was due to Motor-related Cortical Potential (MRCP), which is a super low-frequency negative shift in EEG recording. MRCP may come earlier than the movement because the cortical processes are employed in the planning and preparation of movement. Hence, MRCP can be used for predicting the movement in BCI applications. In this experiment, subjects did not know which movement and when they were going to execute. Hence the MRCP response came out after the zero point. The brain area of its response was the same as the motor area of the corresponding action.
- ROI #2 (from 1 to 3 s, 9 to 13 Hz): The ERD response from the *μ* rhythm oscillation was observed in this region. By carefully comparing the responses of RH and LH, we found that the ERD phenomenon happened in motor areas of both left and right hemispheres. But the ERD in the contralateral brain area was larger than that in the ipsilateral brain area.
- ROI #3 (from 1 to 3 s, 20 to 30 Hz): The ERD response from the *β* rhythm oscillation was also well reported for motor-based BCI application. Similar to ROI #2, the ERD phenomenon happened in the motor areas of both the left and right hemispheres for RH and LH. The interhemispheric difference was mainly located in the parietal-central area in ROI #2, and located in the frontal-central area in ROI #3.
- ROI #4 (from 1 to 3 s, 55 to 90 Hz): During the movement, high *γ* oscillation also showed an interhemispheric difference for RH and LH. But the magnitude was weaker than that of ROI #2 and ROI #3. We found that the magnitude change in ROI #4 was ERD for both RH and LH, which was ERS for FT. But the magnitude was quite small for the high-frequency EEG recording.
- ROI #5 (from 4 to 4.5 s, 12 to 14 Hz): After the movement, *μ* rhythm ERS phenomenon occurred in the corresponding motor area for FT, RH, and RL. With the large sample size in our dataset, some more detailed results can be observed. For example, the *μ* rhythm ERS for FT occurred earlier at a higher frequency band (around 12.5 Hz, and 4.2 s) than ERS for RH and LH (around 9 Hz, and 5 s).
- ROI #6 (from 4 to 4.5 s, 20 to 30 Hz): *β* rhythm ERS phenomenon was seen in this region. We found that from 3.7 to 4.2 s, the *μ* rhythm ERD and *β* rhythm ERS occurred simultaneously for RH and RL. The *μ* rhythm ERS and *β* rhythm ERS did not occur at the same time for RH and LH, but occurred at almost the same time interval for FT.

### 4.2 Benchmark results

#### 4.2.1 Offline evaluation

Since this section only aims to evaluate the performance degradation resulting from larger population sizes, cross-session, and cross-task tests, only the MFCC feature and SVM classifier was adopted. The results for the investigation of the uniqueness of individuals, robustness to mental state changes, and permanence over recording sessions of EEG-based biometrics are shown in Fig. 7(A-C) correspondingly.

**Fig. 7.**
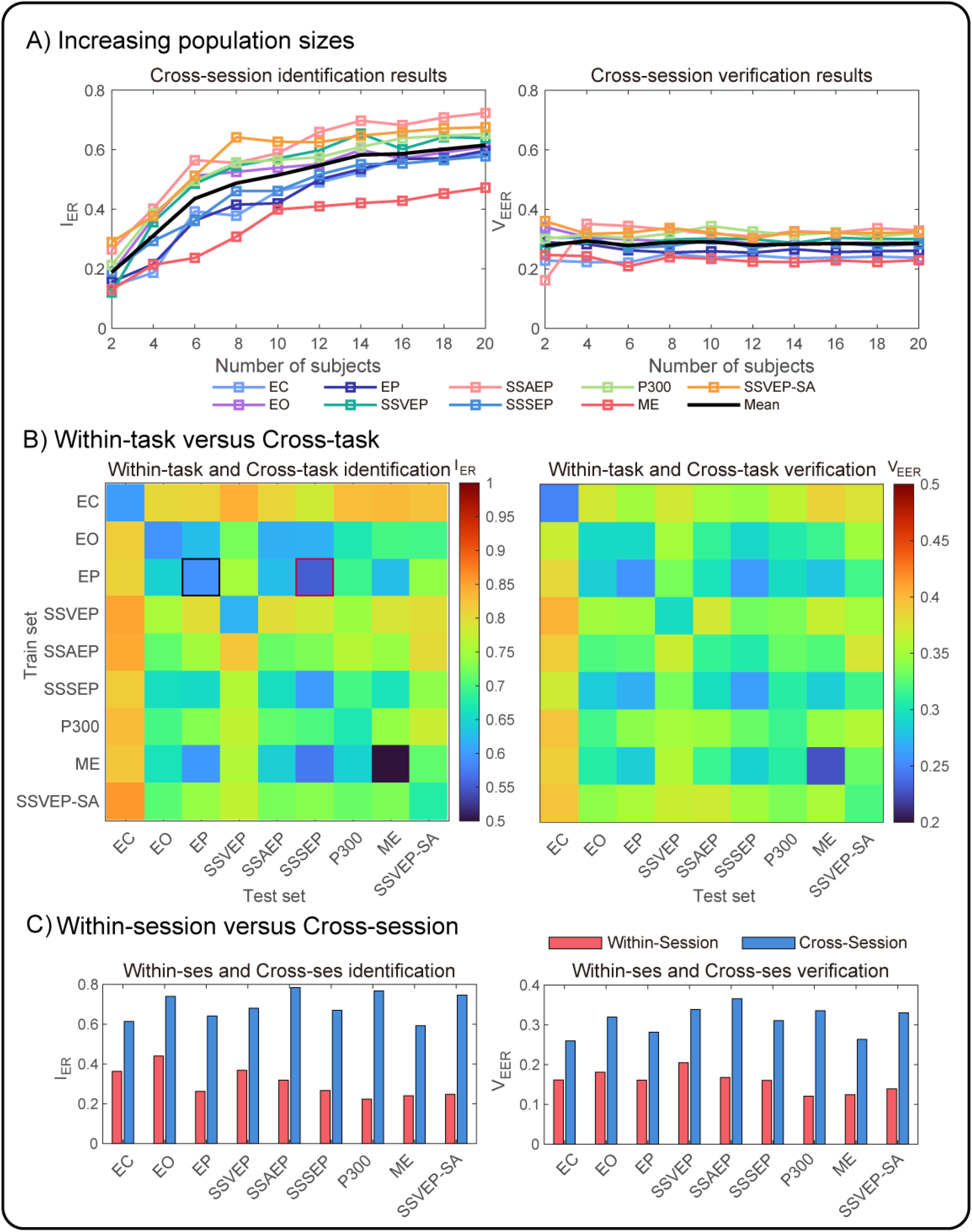
The influence of (A) increasing population sizes, (B) cross-task tests, and (C) cross-session tests on the performance degradation of EEG-based biometric systems. These results were obtained on 20 subjects in the enrollment set and calibration set. *Left:* identification results. *Right*: verification results.

In terms of the uniqueness of individuals, it can be observed that the averaged identification error rate *I*_ER_ increased from 0.81 to 0.34 as the population sizes increased from 2 to 20, as shown in the left figure of Fig. 7A. It should be noted that with the different population sizes, the chance levels are different. With the population sizes rising to a larger scale (e.g., 106 subjects in our testing set), the identification problem for machine learning algorithms to identify one subject would be more challenging. Since the verification task is a one-to-one matching problem, increasing the population size will not lead to its performance degradation.

Fig. 7B showed the robustness of EEG-based biometrics to mental state changes, in which the row represents the task used for training, and the column represents the task used for testing. In the left figure of Fig. 7B, the red square border (where EP was used to train and SSSEP was used to test) shows a lower identification error rate than the black square border (where EP was used to train and test), which is 0.55 versus 0.59. Except for this, the minimal error rate for other rows was achieved at the diagonal line, in which the training task and the testing task is the same. This result is consistent for both identification and verification tasks. Due to the unequal number of training samples for each task, it’s unfair to compare the error rate across different rows. The within-task and cross-task test results were obtained in cross-session settings.

As for the permanence across recording sessions in Fig. 7C, the averaged equal error rate for verification *V*_*EER*_ increased from 0.16 to 0.31, and the average classification error rate for identification *I*_*ER*_ increased from 0.30 to 0.69.

As illustrated in Fig. 8, 6 performance metrics including (*I*_*ER*_, *V*_*ER*_, *V*_*FAR*_, *V*_*FRR*_, *T*_*train*_, and *T*_*test*_) were displayed on radar charts for the six modes for the combination of feature extraction (PSD, AR, MFCC) and classification (L2 and SVM) methods. The smaller area in the radar charts indicates better performance of the biometric system. The value of *TER*, which decides the ranking on the leaderboard for six models, is also provided. PSD+L2 provided the best result as *TER* = 0.278.

In terms of extracted features, the AR features have the worst performance no matter which classifier was used. PSD features have a better or similar performance, as compared with more elaborated features such as MFCC across each metric. It is interesting to note that with the threshold determined by the equal error rate (i.e., *V*_*FAR*_ = *V*_*FRR*_) in an independent training set, the performance of MFCC and AR features are very different on the metrics of *V*_*FAR*_ and *V*_*FRR*_. Especially, the *V*_*FAR*_ of AR feature is 0.628 and 0.578 for L2 and SVM, but the corresponding *V*_*FRR*_ is 0.096 and 0.162, exhibited strong instability for AR-based verification system.

In terms of classifier, the SVM classifier performed a little better than L2 classifier with the PSD and MFCC features. As for the time consumption, since the L2-based classifier has to compare the new input with all points in the training set, its evaluation time *T*_*test*_ is slower than the SVM classifier.

#### 4.2.3 Online Submission

For the example submission, we get the score of 0.4039, which indicated the total error rate *TER* of 59.61% in leaderboard A. It should be noted that the methods with a better result on the simulated online analysis may not necessarily directly lead to higher scores in the online submission. That is caused by the different samples used for training and testing. For the same reason, it is also possible that the score varies from Leaderboard A to Leaderboard B.

**Fig. 8.**
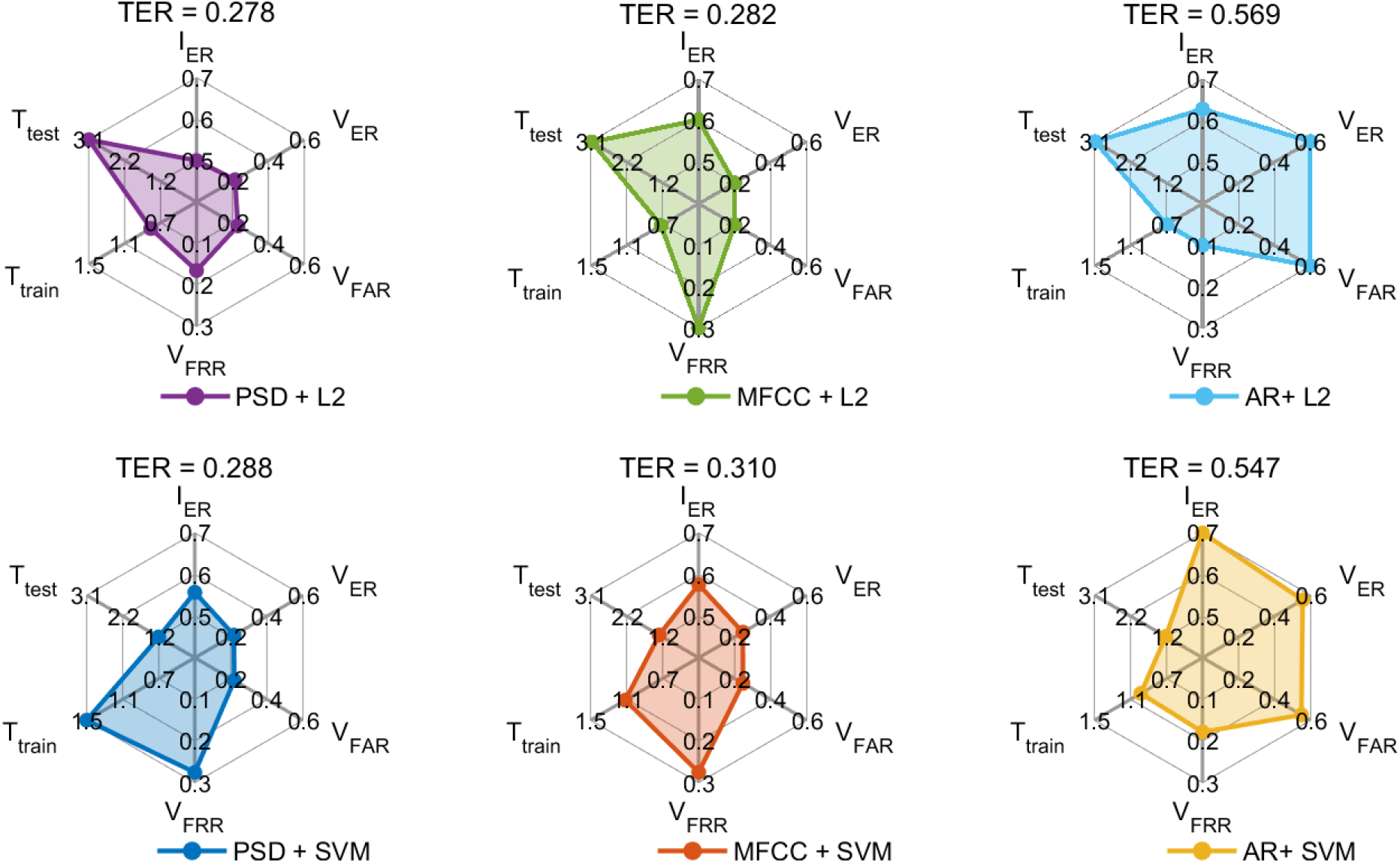
Model evaluation for the combination of different features (PSD, MFCC, SVM) and different classifiers (L2 and SVM). The limit of each ax in the radar plot was decided by the minimal value and the maximal value for each metric among the six models.

## 5. Discussion and Conclusion

### 5.1 Challenges in EEG-based biometrics competition

In the field of EEG-baed biometrics, many studies have mentioned that their major limitation is lacking a database containing large sample sizes, recorded under different conditions and different sessions (Wang et al., 2020; Chan et al., 2018; Delpozo-Banos et al., 2018). Based on our proposed M^3^CV database, we launch the EEG-based biometric competition. In view of the current development of EEG-based biometric technology, this competition proposes the following challenges for machine learning algorithms from many aspects:

1. **Challenge with large population size**: the population size of the testing database is an important indicator of whether the current EEG-based biometric technology can be practically applied. As illustrated in Fig. 7(A), the identification error increases greatly with the increase of the population size. The identification task on the 106 population size would be more challenging for the machine learning algorithm.
2. **Challenge for cross-session tests**: a practical biometric system should be robust to cross-session variability. However, as illustrated in Fig. 7(C), the machine learning model would easily achieve a good performance in the within-session test but degraded severely in the cross-session test. Hence, the cross-session test would be one big challenge in this competition.
3. **Challenge for cross-task tests**: similar to the cross-session tests, the robustness of the biometric system to the cross-task variability is also a challenge in this competition. For the real application, the biometric system should recognize the subject even when they are in different mental states.
4. **Challenge for intruders**: Whether the biometric system can prevent an attack from intruders outside the enrollment set is determined by the identification performed on a closed set or an open set (Kumar et al., 2021). That would lead to very different strategies for the classifier design in the use of machine learning technique

To our best knowledge, no previous study has tested the permanence and robustness of EEG-based biometric features over 100 subjects in an open-set setting. Hence, the M^3^CV database can facilitate the development of advanced machine learning algorithms for EEG-based biometrics.

### 5.2 From competition to real application

Due to the limitations of the competition conditions, the winner of the competition may still be some distance from the real application.

Firstly, the competition needs to have a unique ranking. Hence the score 1 − *TER* was applied to comprehensively evaluate the contestant’s performance in the identification and verification task to decide the final winner. However, the evaluation of a system’s performance should consider multiple aspects. On the calibration set, the performance metrics of *I*_*ER*_, *V*_*ER*_, *V*_*FAR*_ *V*_*FRR*_, *T*_*train*_, and *T*_*test*_ have been proposed for a comprehensive evaluation of the system. For the submission of each team, the error rate under different conditions can be calculated offline, but the computation complexity of training and testing is difficult to make a relatively consistent evaluation. Hence, it is required for the contestants to report the time for training and testing, and their hardware and software environment.

Furthermore, the baseline models provided in the studies are all one-class classifiers (L2 and one-class SVM). If a new subject is added to the enrollment set, the biometrics system just needs to add the model for the new coming subject alone. It doesn’t need to retrain the whole classifier model. We believe the multi-class classifier would achieve better performance in the competition. But one-class classifier would be more practical in the real application.

### 5.3 Intra- and inter-subject variability

Intra- and inter-subject variability poses a great challenge for the interpretation and decoding of EEG signals (Seghier and Price, 2018; Wei et al., 2021). Traditional ERP analysis and statistical test methods were typically used to analyze the group-level commonality of EEG signals, while machine learning techniques are considered to be more powerful in dealing with intra- and inter-subject variability. The dataset for this competition is not only restricted to the study of EEG-based biometric algorithms but can also be used for other applications, such as cross-session reliability analysis of EEG signals from different tasks, and transfer learning in cross-subject and cross-session BCI studies. Furthermore, the cross-run and cross-trial variability within subject is also a meaninful topic to study. The multirun EEG recording of resting-state, Transient-state sensory, motor execution also made the M3CV database useful in these studies.

### 5.4 Deep learning

Undoubtedly, deep learning is the biggest advance in machine learning in recent years, which has also been applied in the EEG-based biometrics study (Behrouzi and Hatzinakos, 2022; Wang et al., 2019; Maiorana, 2021b). However, due to the lack of support from large datasets, the generalizability of deep learning has always been doubtable in this field. On the other hand, most of the newly proposed ideas of transfer learning are based on the framework of deep learning. Based on this competition on the M3CV database, it is hoped that some cross-session and cross-task deep learning methods can emerge. Furthermore, it is also hoped that these methods can not only work well in the field of EEG-based biometrics but also provide a universal solution in dealing with Intra- and inter-subject variability of the EEG signals.

### 5.5 Conclusion

The lack of a large-scale comprehensive EEG database, which contains data recorded from multiple subjects in multiple sessions and over multiple tasks, limits our understanding of intra- and inter-subject variability and hinders the development of advanced machine learning algorithms. In this study, we established an M^3^CV database with 106 subjects, 2 sessions, and 6 tasks to reveal the commonality and variability of cross-subject, cross-session, and cross-task EEG signals. Based on this M^3^CV dataset, a machine learning competition about EEG-based biometrics was launched to help this field grow. Except for advancing the development of machine learning algorithms for EEG decoding (such as biometrics and BCI), we believe the M^3^CV database can help researchers gain a deep understanding of the relationship between different types of EEG signals, such as resting state, motor, sensory-related, or cognitive-related EEG signals.

## Acknowledgements

We thank Hu Li and Kong Yazhuo from the Institute of Psychology, Chinese Academy of Sciences, Peng Weiwei from the School of Psychology, Shenzhen University, Liang Meng from the School of Medical Imaging, Tianjin Medical University, Liu Jixin from the School of Life Science and Technology, Xidian University for their suggestions on our experimental design.

